# Chromosomal Resistance to Metronidazole in *Clostridioides difficile* can be Mediated By Epistasis Between Iron Homeostasis and Oxidoreductases

**DOI:** 10.1101/2020.03.04.977868

**Authors:** Aditi Deshpande, Xiaoqian Wu, Wenwen Huo, Kelli L. Palmer, Julian G. Hurdle

**Author notes:** Correspondence and requests for materials should be addressed to J.G.H. These authors equally contributed (Aditi Deshpande, Xiaoqian Wu). Author order was determined both alphabetically and in order of increasing seniority.

## Abstract

Chromosomal resistance to metronidazole has emerged in clinical *Clostridioides difficile*, but the genetic mechanisms remain unclear. This is further hindered by the inability to generate spontaneous metronidazole-resistant mutants in the lab to aid genetic studies. We therefore constructed a mismatch repair mutator, in non-toxigenic ATCC 700057, to unbiasedly survey the mutational landscape for *de novo* resistance mechanisms. In separate experimental evolutions, the mutator adopted a deterministic path to resistance, with truncation of ferrous iron transporter FeoB1 as a first-step mechanism of low level resistance. Allelic deletion of *feoB1* in ATCC 700057 reduced intracellular iron content, appearing to shift cells toward flavodoxin-mediated oxidoreductase reactions, which are less favorable for metronidazole’s cellular action. Higher level resistance evolved from sequential acquisition of mutations to catalytic domains of pyruvate-ferredoxin oxidoreductase (PFOR encoded by *nifJ*); a synonymous codon change to *xdhA1* (xanthine dehydrogenase subunit A), likely affecting its translation; and lastly, frameshift and point mutations that inactivated the iron-sulfur cluster regulator (IscR). Gene silencing of *nifJ, xdhA1* or *iscR* with catalytically dead Cas9 revealed that resistance involving these genes only occurred when *feoB1* was inactivated i.e. resistance was only seen in an *feoB1*-deletion mutant and not the isogenic wild-type parent. These findings show that metronidazole resistance in *C. difficile* is complex, involving multi-genetic mechanisms that could intersect with iron-dependent metabolic pathways.

## INTRODUCTION

*Clostridioides difficile* infection (CDI) is a leading cause of diarrhea in hospitalized patients in developed countries. Since 2003, the emergence and spread of epidemic strains has significantly increased the incidence and severity of CDI. The health care impact of the epidemic *C. difficile* strains is evident from there being about half a million cases of CDI and ∼29,000 deaths in the United States in 2011 (1, 2).

Owing to its potent anti-anaerobic activity and low cost, metronidazole was traditionally preferred to treat mild to moderate CDI (3). Metronidazole is a 5-nitroimidazole prodrug that is primarily activated in anaerobes conducting reactions that generate low redox potentials (4). Enzymatic reduction of metronidazole occurs via reactions catalyzed by oxidoreductases, such as pyruvate-ferredoxin oxidoreductase (PFOR), and involves the transfer of electrons to its nitro group from cofactors like ferredoxin and flavodoxin. This produces an unstable nitroimidazole anion that may be converted to reactive desnitro, nitroso and hydroxylamine intermediates, which react with DNA, proteins and non-protein thiols to form adducts (4).

The efficacy of metronidazole has declined with the emergence of epidemic strains. Therefore, in the 2017 IDSA/SHEA treatment guidelines, metronidazole is no longer a first-line drug for adult CDI (5). However, there is a need for scientific evidence explaining the decline in metronidazole efficacy, as continued use of the drug is likely until the new guidelines become standard practice. The decline in metronidazole efficacy appears to correlate with the emergence and spread of resistant strains of different ribotype backgrounds (6-8); metronidazole resistance is defined by the EUCAST breakpoint of >2 μg/ml. For example, Snydman reported that ∼8% of U.S. isolates (2011-2016) are resistant to metronidazole (7), while the rate in Europe ranged from 0.1% to 0.5% for isolates collected between 2011 to 2014 (6).

The foremost attempt to characterize the genetic basis for chromosomal metronidazole resistance in a clinical strain was by Lynch *et al*. (9). The patient isolate strain was isolated on metronidazole-containing agar, but was then passaged, *in vitro*, in drug and was found to contain mutations in PFOR, ferric uptake regulator (Fur) and the oxygen-independent coproporphyrinogen III oxidase (HemN) among other changes. In another study, Moura *et al*. evaluated a non-toxigenic metronidazole-resistant *C. difficile* showing it adjusted metabolic pathways that are linked to PFOR activity (10). Most recently, Boekhoud *et al*. (11) reported that *C. difficile* clinical isolates carried a conjugative plasmid (pCD-METRO) conferring resistance to metronidazole, but the plasmid was not present in several CDI associated ribotypes that were metronidazole-resistant. This indicates that pCD-METRO is not a universal mechanism of resistance; furthermore, the genetic determinants on pCD-METRO directly responsible for the phenotype are still unknown. Taken together, it appears *C. difficile* could have multiple ways to evolve metronidazole resistance, but these mechanisms remain unclear or require genetic and biochemical validation in naïve hosts. In contrast, mechanisms of MTZ resistance in several pathogens for which the drug is prescribed are known e.g. *Helicobacter pylori* and *Entamoeba* histolytica (4).

There have been two critical barriers to knowledge of metronidazole resistance in *C. difficile*. Firstly, metronidazole-resistant mutants are designated to be unstable, as evident by inconsistent metronidazole susceptibility profiles of the strains (9, 12). Secondly, there is an inability to select laboratory mutants to allow for controlled genetic studies (13). To address these challenges, from a laboratory perspective, we constructed a mutator tool by deleting the DNA mismatch repair system in a non-toxigenic *C. difficile* strain. Mutators are employed for accelerated evolution to study mutation accumulation in bacteria (14, 15). Using this concept, we investigated the genomic landscape of *C. difficile* for *de novo* resistance mechanisms that are evolutionarily feasible, albeit *in vitro*. Hence, we obtained new insights that *C. difficile* can manipulate iron uptake and iron-dependent oxidoreductive pathways to develop resistance to metronidazole.

## RESULTS AND DISCUSSION

### Hypermutator construction by deleting DNA mismatch repair (MMR) genes

The MutSL MMR proteins are involved in correcting replicative errors (16), wherein MutS identifies mispaired or unpaired bases and recruits the endonuclease MutL that initiates the removal and repair of misincorporated bases (16). *C. difficile* was found to carry adjacent *mutS* and *mutL* genes (CD630_19770 and CD630_19760), in addition to the *mutS* homolog *mutS2* (CD630_07090). To construct a *C. difficile* mutator, we chose the strain ATCC 700057 (17) because it is non-pathogenic, lacking the toxin genes *tcdA* and *tcdB* (18), and is widely used for antibiotic susceptibility testing (12). We individually deleted *mutS, mutL, mutS2* and the entire *mutSL* operon by allelic exchange using pMTL-SC7215; deletions were confirmed by PCR (19). Mutability was assessed from the frequency of isolating mutants to the antibiotics rifaximin and fidaxomicin at 4 x their MICs (0.5 and 0.25 μg/ml, respectively). As shown in **Fig. 1A** mutation frequencies were enhanced upon deletion of *mutS* or *mutL*, but *mutS2* had no effect. Deletion of *mutSL* caused the highest increase (>80-fold) in mutability with respect to WT ATCC 700057 i.e. mutation frequencies were 10^−6^ for 700057Δ*mutSL* versus 10^-8^ for the WT ATCC 700057; and this strain was significantly more mutable than Δ*mutS* or Δ*mutL* strains, as determined by unpaired t-tests (**Table S1**). Mutability was reversed when 700057Δ*mutSL* was complemented with WT *mutSL* (**Fig. S1**). Thus, 700057Δ*mutSL* (i.e. mutator) was used to select metronidazole-resistant mutants below.

**Figure 1.**
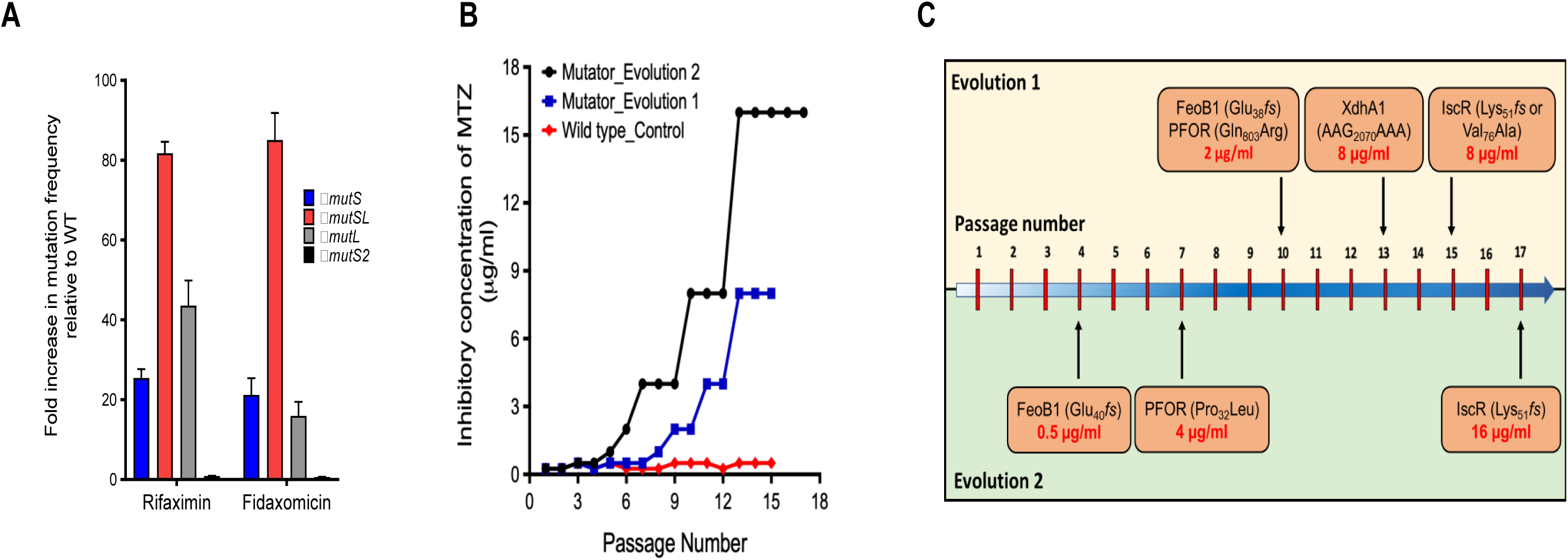
Evolution of metronidazole (MTZ) resistance using mismatch repair (MMR) deficient mutator. (**A**) Rifaximin and fidaxomicin mutation frequencies (MFs) of strains with deletions in MMR genes; fold change in MFs are relative to WT ATCC 700057. Plotted are means ± SEM from three biological replicates. (**B**) Two independent experimental evolutions with the mutator (Δ*mutSL*) resulted in MTZ-resistant mutants, in contrast to the WT. (**C**) The order in which mutation were accumulated in the two evolutions suggest there was a deterministic path to resistance.

### Experimental evolution of metronidazole resistance

The agar MICs of metronidazole against the WT and mutator was 0.25 μg/ml. However, spontaneous mutants could not be selected by plating >10 overnight cultures of each strain onto agars containing 2 and 4 x MICs (*data not shown*). Both strains were therefore serially passaged on agars containing varying concentrations of metronidazole (0.125-16 μg/ml); each passage was incubated up to 3 days to obtain growth. In the first experimental evolution, by the 9^th^ passage, the mutator evolved resistance as its population was inhibited by 2 μg/ml (**Fig. 1B**). By the 15^th^ passage, the population was inhibited at 8 μg/ml. In contrast, stable mutants did not arise from the WT ATCC 700057, even up to 15 serial passages in drug (**Fig. 1B**) and is consistent with prior reports of an inability to isolate *in vitro* metronidazole-resistant mutants of *C. difficile* (13). To further comprehend the evolutionary path to resistance, we conducted a separate experimental evolution with the mutator and identified that by the 6^th^ and 17^th^ passages the population was inhibited by 2 and 16 μg/ml of drug, respectively (**Fig. 1B, C**).

### Identification of genetic changes associated with *de novo* metronidazole resistance

As expected, the mutator accumulated insertions, substitutions and deletions across the genome and showed no genomic site specificity (**Fig. S2**). Functional gene classification showed that mutations occurred to iron transporters, iron-sulfur proteins, oxidoreductases, carbohydrate metabolism, fatty acid metabolism, cell surface/division proteins and other genes (**Fig. S3**). However, to study mutations associated with resistance, we focused on proteins involved in cellular redox and metal homeostasis that may affect the activation of metronidazole.

#### (i) Identification of genetic changes in evolution 1

From the culture population at the endpoint of the evolution experiment 1 (passage 15), we isolated and genome sequenced three mutant colonies, designated as JWD-1 (MIC=8 μg/ml), JWD-2 (MIC=32 μg/ml) and JWD-3 (MIC=32 μg/ml). They all carried a frameshifting deletion (Glu38*fs*) in the FeoB1, encoded by CD630_14790; a substitution in PFOR (Gln_803_Arg) encoded by *nifJ* (CD630_26820); and a synonymous change (AA**G**_2070_AA**A**) in xanthine dehydrogenase subunit *xdhA1*, encoded by CD630_20730. JWD-2 and JWD-3 also carried unique changes of Lys_51_*fs* and Val_76_Ala respectively in the iron-sulfur cluster regulator (IscR), encoded by CD630_12780. These mutations in IscR may explain why JWD-2 and JWD-3 were 4-fold more resistant to metronidazole than JWD-1, which lacked changes to the regulator. We identified the order in which mutations arose (**Fig. 1C**) by determining the MICs and Sanger sequencing of three individual mutant colonies each from passages 10 and 13. Mutants from passage 10 had MICs of 2 μg/ml and carried the above variations in PFOR and FeoB1, while mutants (MICs of 8 μg/ml) from passage 13 also carried the synonymous change in *xdhA1.*

#### (ii) Identification of genetic changes in evolution 2

JWD-4 (MIC=64 μg/ml), an endpoint mutant (passage 17) of experiment 2 was isolated and genome sequenced. Similar to the above endpoint mutants, JWD-4 contained disruptions to FeoB1 (Glu_40_*fs*), PFOR (Pro_32_Leu) and IscR (Lys_51_*fs*), but also acquired a Gly_270_Asp substitution in xanthine permease (CD630_20910). The order in which mutations arose (**Fig. 1C**) were also determined by MIC testing and Sanger sequencing of minimum three colonies per time point and metronidazole MICS were also measured. This revealed FeoB1 was disrupted by passage 4 (MIC=1 μg/ml); PFOR by passage 7 (MIC=2 μg/ml); xanthine permease by passage 14 (MIC=16 μg/ml); and IscR by passage 17 (MIC=64 μg/ml). The occurrence of similar genetic changes in the two independent serial passage experiments suggests there was a deterministic path to metronidazole resistance.

### Explanation of *de novo* genetic variations associated with metronidazole resistance

#### (i) FeoB1 participation

Under anaerobic conditions, iron mostly exists in ferrous (Fe^2+^) form. In *C. difficile*, FeoB1 is predicted to be the main iron transporter, as it is the most upregulated iron transporter in low iron conditions and *in vivo* in hamsters (20, 21). *In vitro*, the homologs FeoB2 and FeoB3 are thought to be less responsive to changes in iron (20). Thus, the loss of FeoB1 may have lowered the supply of iron to iron carrier proteins that mediate electron transfer to metronidazole (22). In support of our findings, deletion of *feoAB* in *B. fragilis* conferred a 10-fold decrease in metronidazole activity (23).

#### (ii) Oxidoreductases PFOR and XDH participation

PFOR catalyzes the oxidation of pyruvate to acetyl-CoA, while XdhA1 is the molybdenum-binding subunit of xanthine dehydrogenase (XDH) that catalyzes the oxidation of purines. Sequence alignment of PFOR homologs of *C. difficile* (CD630_26820) *and Desulfovibrio africanus* (Desaf_2186) (59.4% similarity) indicated that the Pro_32_Leu substitution occurred adjacent to the critical threonine-31 site that forms the catalytic domain for pyruvate binding, whereas Gln_803_Arg occurred in domain VI upstream of cysteine-815 that binds the proximal [4Fe-4S] cluster (24, 25) (**Fig. S4**). These substitutions likely affected the enzyme’s catalytic activity, involving electron transfer to electron carrier proteins. To evaluate the synonymous change in *xdhA1*, we analyzed its secondary structure using the RNA folding algorithm mfold (26) to gauge potential effects on RNA translation. The AA**G**_2070_AA**A** change (**Fig. S5**) was predicted to introduce an unstable stem loop, with a positive free energy change (ΔG = 3.8 kcal/mol for the mutant and ΔG = −25.9 kcal/mol for the WT), which is unfavorable for RNA translation (27). With regard to xanthine permease, the role of this transporter in metronidazole resistance is presently unclear, although we speculate that Gly_270_Asp substitution might affect the supply of xanthine to XDH.

#### (iii) IscR participation

Iron-sulfur [Fe–S] clusters are essential to the biochemistry of several proteins that conduct electron transfer reactions (28, 29). In pathogenic bacteria the assembly and incorporation of [Fe–S] clusters is mostly controlled by the *isc* and *suf* operons, of which *isc* is the house-keeping system; furthermore, *C. difficile* lacks the *suf* operon (30). Holo-IscR binds DNA as a homodimer, but damage to its [4Fe-4S] by reactive free radicals or iron-starvation increases cellular levels of apo-IscR, triggering defensive mechanisms including antioxidants such as cysteine and non-protein thiols (31). The Lys51*fs* variation, in strains JWD-2 and JWD-4, disabled the function of IscR, by forming an aberrant protein lacking cysteines-92, 98 and 104 that are required to bind [4Fe-4S] (32). To characterize the effect of the Val_76_Ala substitution in IscR in JWD-3, we analyzed the crystal structure of dimeric IscR bound to DNA from *E. coli* (pdb ID: 4CHU) using UCSF Chimera (33). The IscRs from *C. difficile* ATCC 700057 and *E. coli* K12 are closely related with ∼41% identical amino acids and valine-76 is a conserved amino acid. In the dimeric structure (**Fig. S6**), valine-76 on Chain A occurs with other lipophilic residues in a dimer interface where it hydrophobically interacts with leucine-113 on Chain B. It is plausible that the less hydrophobic alanine-76 reduces hydrophobic steric packing for stabilizing the dimeric DNA binding conformation of IscR.

### FeoB1-mediated cellular changes associated with metronidazole resistance

#### (i) *Deleting feoB1* affects resistance and iron-content

Since *feoB1* was inactivated early in the serial evolution (**Fig. 1C**), we first examined the role of this gene by deleting it in ATCC 700057. The deletion mutant (700057*ΔfeoB1*) grew in 0.5 μg/ml of metronidazole (MIC=1 μg/ml), whereas the WT strain grew in 0.125 μg/ml (MIC=0.25 μg/ml) of drug (**Fig. 2A**). Susceptibility was restored by complementing the *ΔfeoB1* strain with WT *feoB1*, expressed from its own promoter in pMTL84151 (**Fig. 2A**). There was ∼21% lesser iron content in the mutant (0.0753 ± 0.002 ppm) compared to the WT (0.0956 ± 0.004 ppm), as determined by ICP-OES. Consistent with reduced intracellular iron content, 700057Δ*feoB1* also showed increased transcription of *fur* (3.9 ± 0.17-fold) and ferrichrome ABC transporter subunit (*fhuB*) (5.13 ± 0.25-fold), compared to the WT, in the absence of drug (**Fig. 2B**). In drug, the Δ*feoB1* mutant showed 10.50 ± 0.96 (*fur*) and 9.16 ± 0.25 (*fhuB*) fold higher gene expression than the WT (**Fig. 2B**).

**Figure 2.**
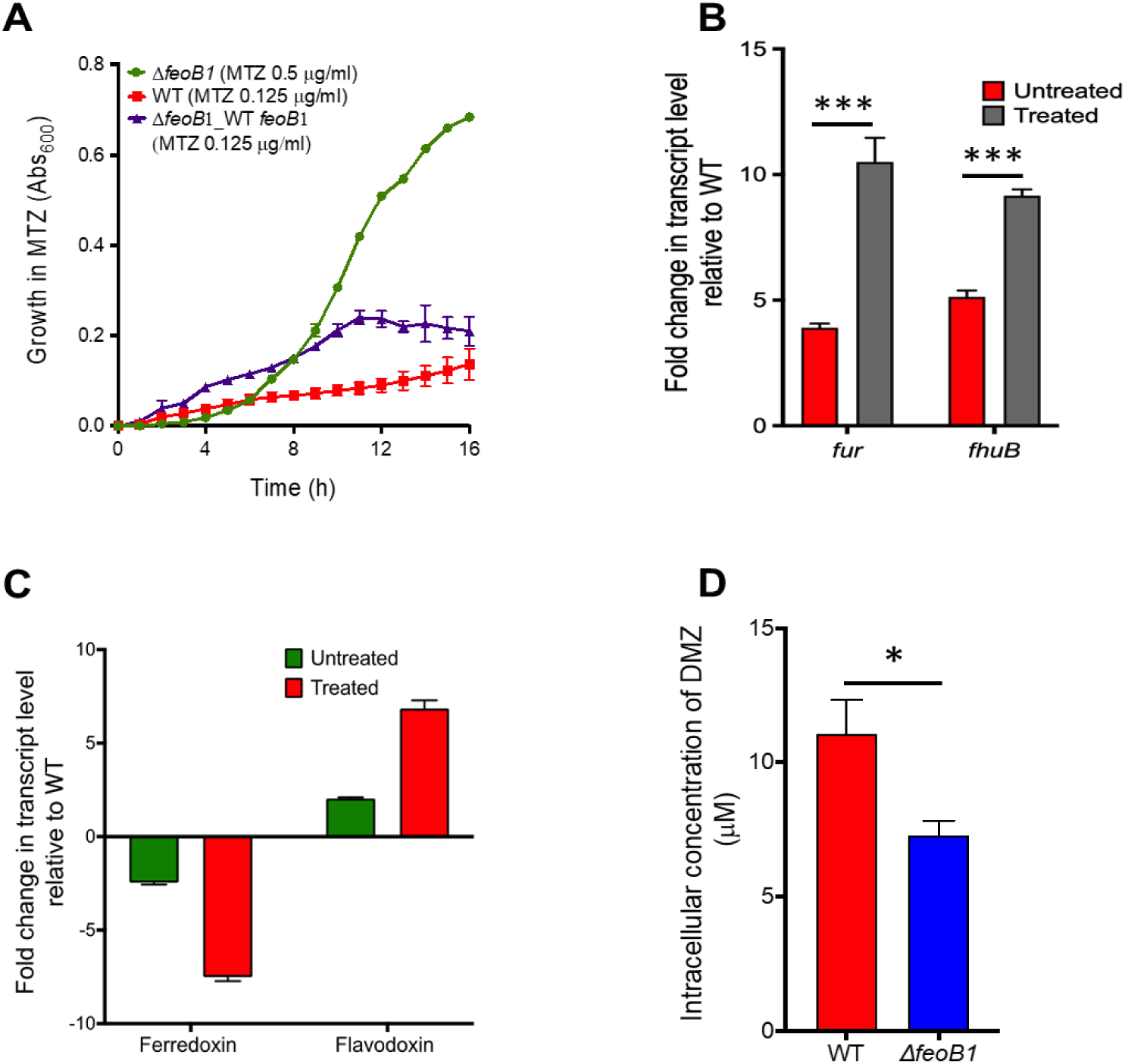
Effect of *feoB1* deletion on: (**A**) growth in metronidazole (MTZ). The Δ*feoB1* mutant grew better in MTZ, as shown from optical density readings [Abs_600_nm]. MICs were 1 µg/ml against the Δ*feoB1* mutant and MIC=0.25 µg/ml against the WT and complemented Δ*feoB1* strains. (**B, C**) Transcription of iron-response genes with respect to WT strain, in the absence and presence of MTZ at 4 x MIC against each strain. Increased expression of *fur* and *fhuB* without drug supports iron limitation in Δ*feoB1* mutant, while expression of flavodoxin over ferredoxin suggests a switch to flavodoxin-mediated metabolism. (**D**) Intracellular concentration of dimetridazole (DMZ), after 1 hour, in the Δ*feoB1* and WT strains, suggests less drug accumulation in the mutant.

#### (ii) Effect on expression of cofactors that activate metronidazole

Since the loss of *feoB1* reduced iron content in cells, we reasoned this may diminish metronidazole activation by limiting iron-dependent electron carrier proteins such as ferredoxins. Ferredoxin and flavodoxin are small redox proteins that accept and transfer electrons to metronidazole from oxidoreductases, such as PFOR. Transcriptional analysis revealed that in the absence of drug, 700057Δ*feoB1* downregulated (−2.46 ± 0.09-fold) the ferredoxin (*fdx* i.e. CD630_06271) and upregulated (2.04 ± 0.06-fold) the flavodoxin (*fldx* i.e. CD630_19990) relative to WT (**Fig. 2C**). When 700057Δ*feoB1* was exposed to metronidazole, there was further downregulation of *fdx* (−7.51 ± 0.21-fold) and upregulation of *fldx* (6.860 ± 0.43-fold) (**Fig. 2C**). Other ferredoxin and flavodoxin homologs (**Fig. S7**) were lesser expressed supporting published studies that CD630_06271 and CD630_19990 are the most responsive homologs of ferredoxin and flavodoxin, during iron stress in *C. difficile* (20, 21). Ferredoxin is better at transferring electrons to metronidazole, due to carriage of an iron-sulfur cluster that has a lower redox potential than flavodoxin, which carries flavin mononucleotide (34). Thus, these results indicate that loss of *feoB1* led to reduced iron content, prompting a shift to flavodoxin-mediated oxidoreductase reactions that enable cells to resist metronidazole.

#### (iii) Effect on cellular accumulation of nitroimidazole

Since the above results suggest the loss of *feoB1* might decrease metronidazole activation, we quantified the intracellular concentration of dimetridazole, a related nitroimidazole (35). Dimetridazole was used due to the commercial availability of 5-amino-1,2-dimethylimidazole, a likely end-product of dimetridazole activation (35) that could be used as a standard in the LC-MS/MS. Against dimetridazole, 700057Δ*feoB1* showed the same level of resistance to metronidazole (i.e. MIC=1 μg/ml). Concentrated cells were exposed to 1 mM of dimetridazole for 1 hour and intracellular concentrations measured in cell lysates. In 700057Δ*feoB1*, there was 7.31 ± 0.53 μM of dimetridazole (**Fig. 2D**), which represents about 34% lower intracellular drug accumulation, compared to the WT (11.07 ± 1.26 μM). The accumulation of metronidazole and other nitroimidazoles into bacteria is thought to occur in a concentration-dependent manner, whereby as the drug is activated in cells more drug is taken up (36). Thus, the observed lower intracellular accumulation of dimetridazole in 700057Δ*feoB1* might indicate lesser drug activation, which could be expected with a shift to flavodoxin-mediated metabolism. However, we could not detect the amine end-product, which would have provided a measure of drug activation.

### Resistance mediated by *nifJ, xdhA1* and *iscR* appear to require the loss of *feoB1*

To assess the roles played by *nifJ, xdhA1* and *iscR* in metronidazole resistance, we silenced these genes using a xylose-inducible CRISPR-interference vector. In this approach, fusion of antisense nucleotides to the guide RNA of Cas9 blocks gene transcription (37) and is a more facile approach than gene deletions in *C. difficile*. As a positive control, an antisense to *feoB1* was included to measure the effect of gene silencing on growth in metronidazole. Even though gene silencing reduced transcript levels by ∼10-fold (**Fig. 3A**), resistance was only detected in the strain expressing the *feoB1* antisense (MIC = 1 μg/ml); the strain carrying the empty vector was inhibited by 0.25 μg/ml (**Fig. 3B**). Given that *feoB1* mutants arose early in the passages (**Fig. 1C**, we therefore questioned whether expression of resistance involving *nifJ, xdhA1* or *iscR* required the loss of *feoB1*. As shown in **Fig. 3C**, the abovementioned antisenses enhanced survival of 700057Δ*feoB1*, shifting MICs to 8 μg/ml. Thus, the loss of *feoB1* early in the serial evolution, might have influenced the direction of resistance evolution toward genes that when disrupted took advantage of lower iron content (i.e. combinatorial or epistatic resistance).

**Figure 3.**
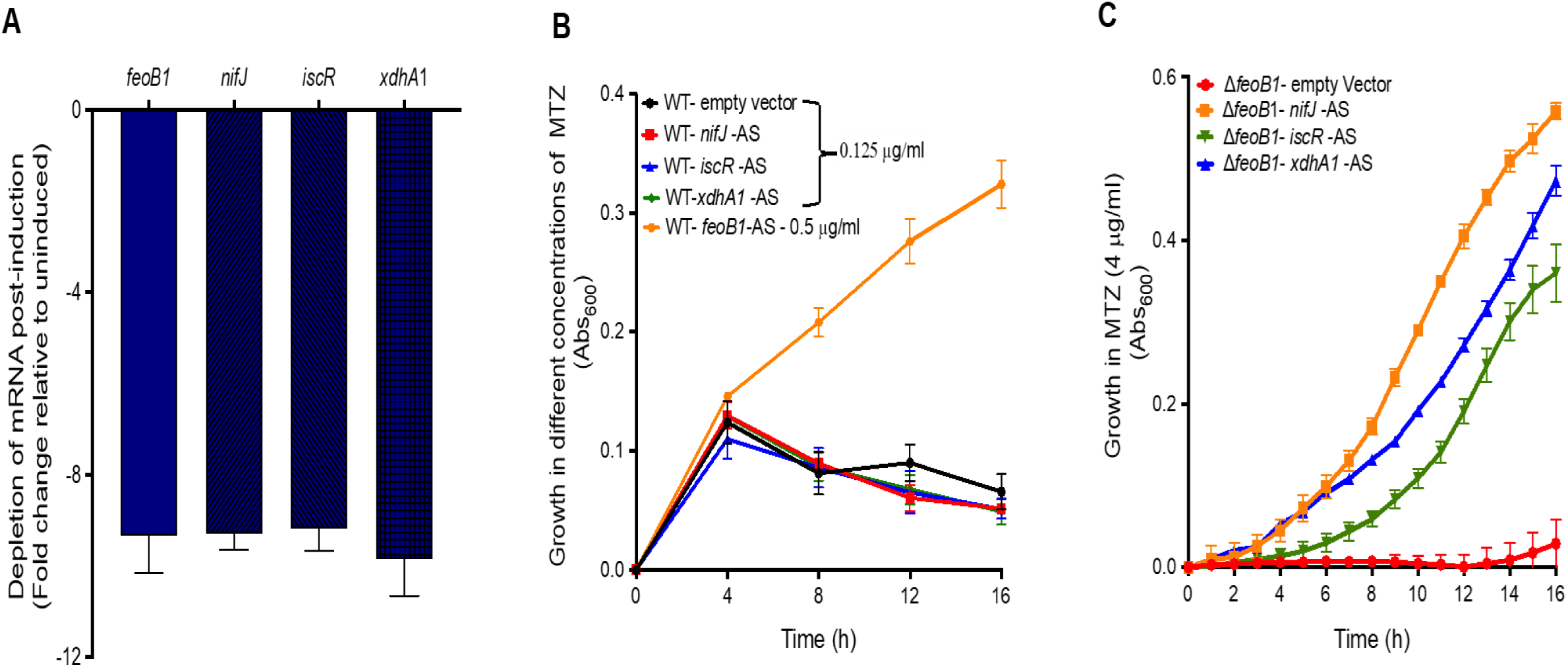
Effect of silencing *nifJ, iscR* or *xdhA1* on resistance in the Δ*feoB1* mutant and WT strain. (**A**) In WT strain depletion of mRNA for *feoB1, nifJ, iscR* and *xdhA1* occur when antisenses are induced with xylose (2% w/v); but (**B**) resistance to metronidazole is seen with only depletion of *feoB1* (MIC=1 μg/ml). (**C**) Conversely, in the Δ*feoB1* mutant, expression of either of the three antisenses caused resistance (i.e. MICs of 8 μg/ml versus 1 μg/ml against Δ*feoB1* mutant).

To confirm that combinatorial resistance was more protective, we exposed cells to 8 μg/ml of metronidazole and analyzed effects on DNA integrity, non-protein thiols and transcription of redox-responsive genes (38, 39). The combinatorial-resistant strains showed more intact DNA, while damage was seen to the DNA of 700057Δ*feoB1*, but was more marked in WT strain (**Fig. 4A**). Likewise, there were higher amounts of free-non-protein thiols in combinatorial-resistant strains (**Fig. 4B**), suggesting a decline in formation of covalent adducts with activated metronidazole. Combinatorial-resistant strains also showed reduced transcription of the DNA damage response gene *recA* (CD630_13280); and redox stress responsive genes (hybrid cluster protein, CD630_21680 and thioredoxin A, CD630_30330) (**Fig. 4C**). Taken together, these results confirm the role of *feoB1* in the evolution of a novel epistatic mechanism of metronidazole resistance. Epistatic mechanisms of antimicrobial resistance are also seen in fluoroquinolone*-*resistant *E. coli*, with double mutations to gyrase A and topoisomerase IV, which produce high-level resistance that is greater than predicted from individual mutations (40).

**Figure 4.**
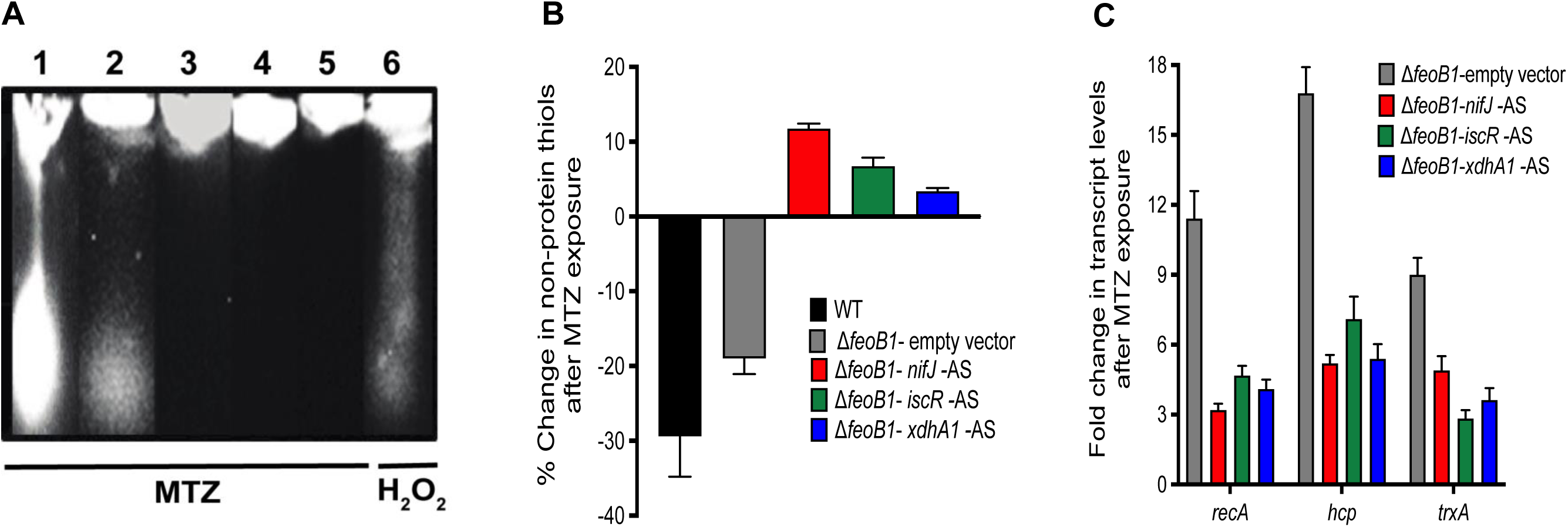
Biochemical and transcriptional validation that higher-level resistance (MIC=8 μg/ml) occurs when mRNA for *nifJ, iscR* or *xdhA1* is depleted in the Δ*feoB1* mutant. (**A**) Metronidazole (MTZ) induced DNA damage was attenuated in the Δ*feoB1* mutant following expression of above antisenses, when compared to Δ*feoB1* and WT strains carrying empty vectors. As a positive control, the WT was exposed to hydrogen peroxide (0.5% v/v) to visualize DNA fragmentation. The image is representative of 3 biological replicates. Resistance was confirmed from there being (**B**) less depletion of non-protein thiols, which form adducts with MTZ; and (**C**) reduced expression of *recA, hcp* and *trxA* that respond to MTZ activity.

### Further assessment of the function of IscR in resistance

As IscR has not been previously linked to metronidazole resistance, we further investigated its role by focusing on pyruvate-lactate metabolism that is affected by iron-sulfur metabolism and is indicative of metronidazole resistance in *B. fragilis* (41). Depletion of *iscR* mRNA in 700057Δ*feoB1* further diminished the transcription of *nifJ* (−6.17 ± 0.29-fold), which was already marginally reduced (−2.67 ± 0.18-fold) with the loss *of feoB1* in 700057Δ*feoB1* compared to WT (**Fig. 5A**). The strain bearing the *iscR* antisense also showed increased levels of pyruvate (1.33 ± 0.03 nM) and lactate (1.86 ± 0.08 nM) compared the parent 700057Δ*feoB1* (0.98 ± 0.02 nM and 1.13 ± 0.045 nM respectively) (**Fig. 5B**). This can be expected if downregulation of *nifJ* further reduced the levels of PFOR protein in cells, which would concomitantly decrease metronidazole activation by the enzyme. Furthermore, a reduction in PFOR activity may also drive the conversion of pyruvate to lactate via lactate dehydrogenase. Therefore, it appears that loss of *iscR* at the endpoint of the evolution (**Fig. 1C**) affected iron-sulfur dependent electron transfer reactions and synergized with both the loss of *feoB1* and mutated oxidoreductase enzymes, which occurred at the earlier evolutionary steps. To confirm the hypothesis that loss of *iscR* affected electron transfer reactions, we tested the susceptibility of cells to the electron acceptor plumbagin, a naphthoquinone that is reduced in reactions catalyzed by oxidoreductases (42). In *Xanthomonas campestris*, the loss of iscR led to plumbagin resistance (43). Our results show that WT ATCC 700057 was inhibited by 0.5 µg/ml of plumbagin, while moderate resistance was seen in 700057Δ*feoB1* (MIC=2 µg/ml). Depletion of *iscR* mRNA in 700057Δ*feoB1* further elevated resistance to plumbagin (MIC=8 µg/ml), in a manner analogous to above observations for metronidazole (**Fig. 5C**).

**Figure 5.**
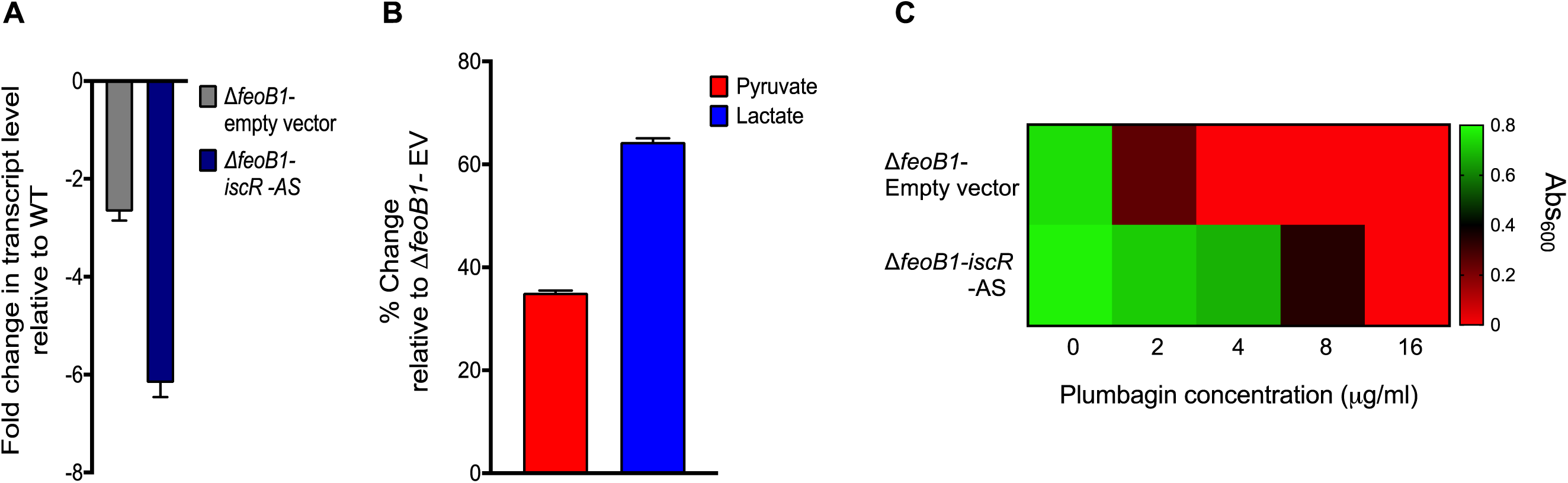
Analysis of how *iscR* inactivation promotes metronidazole (MTZ) resistance in the Δ*feoB1* mutant. (**A**) Depletion of *iscR* mRNA in the Δ*feoB1* mutant caused further downregulation of *nifJ*, which led to (**B**) accumulation of pyruvate and lactate (all values are relative to Δ*feoB1*-empty vector). **(C)** Cross-resistance to plumbagin was seen; the heat-map shows that *iscR* depletion enhances growth in plumbagin (like MTZ, plumbagin is an electron acceptor). These results suggest that loss of *iscR* globally affects iron-dependent electron transfer metabolic reactions, including pyruvate-ferredoxin oxidoreductase.

## CONCLUSION

The involvement of mutations to multiple genes might partly explain why *C. difficile* was slow to evolve chromosomal resistance to metronidazole, in spite of the drug being used since 1980s. While mutations to *feoB1* and oxidoreductases are known to individually cause metronidazole resistance in other organisms (4), to the best of our knowledge there are no reports of epistatic interactions between these mechanisms. Nonetheless, our results generated a model (**Fig. 6**) of metronidazole resistance that could help interpret genetic changes seen in *C. difficile* clinical isolates, which lack pCD-METRO (11). For example, our model in **Fig. 6** generally explains the mechanism of mutations seen a metronidazole-resistant NAP1 *C. difficile* (9, 44); in that study, the strain had mutations to Fur and PFOR (Gly_423_Glu). Disruption of Fur might have affected iron homeostasis to reduce drug susceptibility, which is also shown in *H. pylori* (45). Similarly, the amino acid change in PFOR occurred in domain III, an important subunit for binding of the coenzyme A substrate (25). However, we recognize that FeoB1 is essential for colonization and virulence in other bacteria, which makes it improbable that *C. difficile* will inactivate *feoB1 in vivo* during metronidazole therapy. This raises an important question of whether there are lower free iron concentrations in the gastrointestinal tract that may provide a milieu for resistance to be exhibited by strains with mutated oxidoreductases such as PFOR. Conversely, such resistant strains could be difficult to detect, as current susceptibility testing methods for *C. difficile* use iron-rich medium, and may be categorized as susceptible. Indeed, ineffective detection of metronidazole-resistant *C. difficile* has also been a major concern (9, 12).

**Figure 6.**
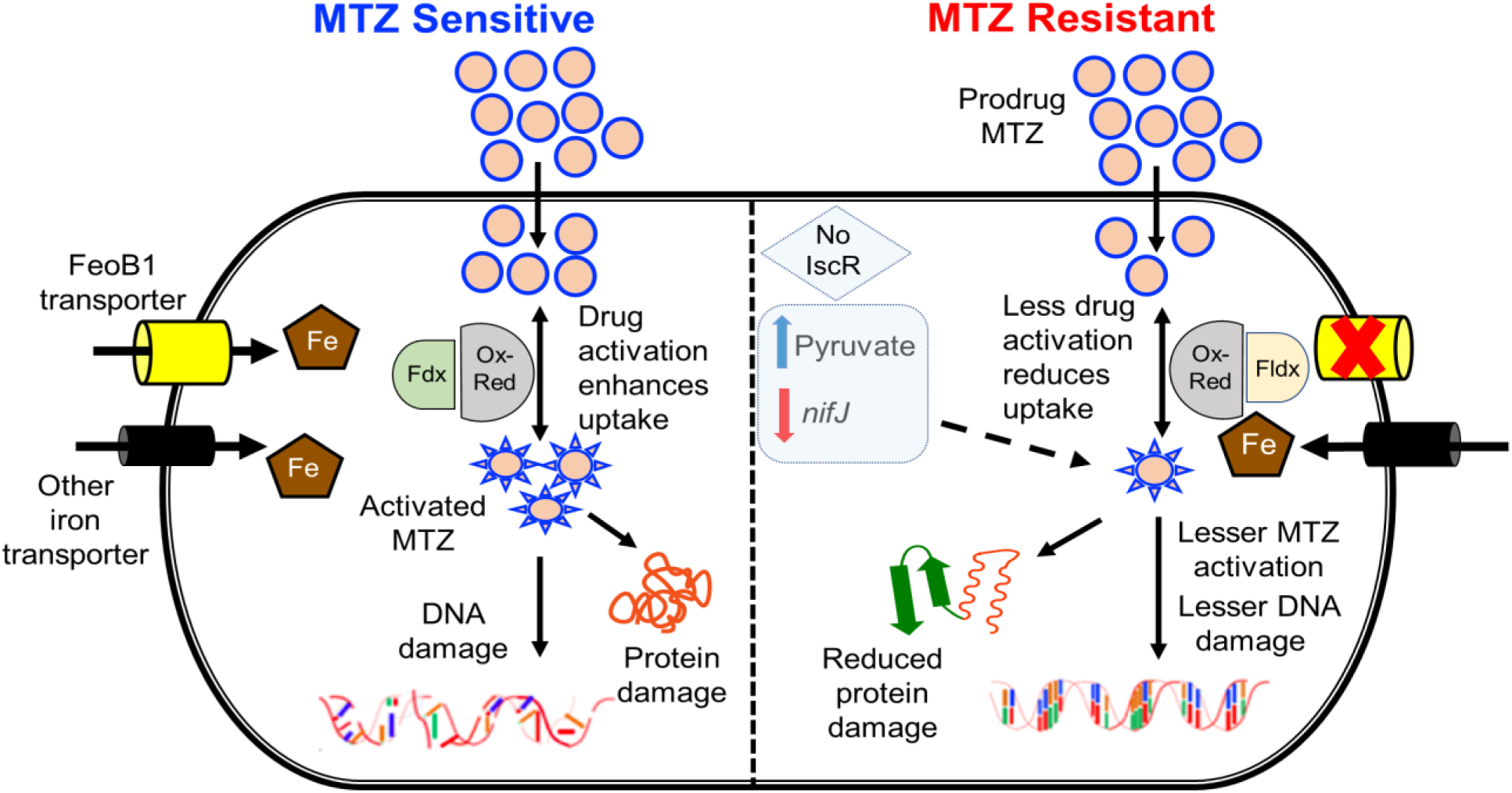
Proposed model of metronidazole (MTZ) resistance in *C. difficile*. (**Left**) In sensitive cells, MTZ is activated by oxidoreductases (Ox-Red; e.g. pyruvate-ferredoxin oxidoreductase [PFOR, *nifJ*]), resulting in free radical damage to DNA and protein damage. Activation then influences drug uptake. Susceptibility is influenced by iron uptake via transporters, mainly FeoB1 (the main iron transporter); and electron carrier proteins ferredoxin (Fdx) with low redox potential. (**Right**) In MTZ-resistant cells, loss of FeoB1 decreases intracellular iron content, probably shifting cells from ferredoxin (Fdx) to flavodoxin (Fldx) mediated metabolism. Fldx is less effective in activating MTZ. Loss of *iscR* synergizes with defective *feoB1* to further reduce metabolic activities that activates MTZ.

A number of outstanding questions remain concerning the apparent multi-genetic mechanisms of metronidazole resistance. Certainly, it is important to establish which resistance mechanisms do or do not impose physiological costs, as this factor might influence the future clinical prevalence of strains. There is also a need to identify whether there are metronidazole-resistant *C. difficile* that also resist oxidative stress, as is known in metronidazole-resistant *H. pylori* and *E.* histolytica that upregulate superoxide dismutase (46, 47). Therefore, in the aftermath of metronidazole use for adult CDI, studies are still warranted to assess the extent to which metronidazole resistance could shape *C. difficile* evolution, epidemiology and pathophysiology.

## MATERIALS AND METHODS

### Strains and culture conditions

Non-toxigenic *C. difficile* ATCC 700057 and derivative strains were routinely grown in pre-reduced Brain Heart Infusion (BHI) broth or agar at 37°C in a Whitley A35 anaerobic workstation (Don Whitley Scientific). *Escherichia coli* strains NEB 5-alpha, CA434 and SD46 were grown at 37°C in LB broth or agar. All strains and plasmids used in this study are listed in **Table S2**. D-cycloserine (250 μg/ml), cefoxitin (8 μg/ml), and thiamphenicol (15 μg/ml) were used to selectively culture *C. difficile* containing plasmids, whereas chloramphenicol (15 μg/ml) and ampicillin (50 μg/ml) or kanamycin (50 μg/ml) were used to grow *E. coli* SD46 and CA4343 respectively. Wherever needed, data was normalized to the protein content.

### Antimicrobial susceptibility

Agar MICs were determined on BHI agar supplemented with hemin (5 μg/ml) and used an inocula of 10^5^ CFU/ml. Agars contained doubling dilutions of compound from 0.06 to 64 μg/ml.

### Genetic manipulation of *C. difficile*

The vector pMTL-SC7215 was used to delete target genes in *C. difficile* by allelic exchange (19). Allelic exchange vectors to delete *mutS, mutS2, mutSL, mutL* and *feoB1* were conjugated into *C. difficile* via *E. coli* donor strains (CA434 or SD46). Allelic exchange was conducted as described (19). Successful gene knockouts were confirmed by PCR. For complementation, *feoB1* was synthesized and cloned into XmaI and XbaI sites of the vector pMTL-84151 by Genscript. Primers used in the study are in **Table S3**. The designed allelic cassettes used in this study are described in the supplementary **Table S4.**

### Gene knockdown

The xylose-inducible vector pXWpxyl-*dcas9* was used to silence the transcription of *feoB1, nifJ, iscR* and *xdhA1*, as previously described (37). The sgRNA (**Table S5**) targeting the abovementioned genes were synthesized and cloned into the vector’s PmeI site by Genscript Biotech (New Jersey). Antisenses were induced with 2% w/v of xylose.

### Determination of mutation frequencies

Briefly, *C. difficile* strains were grown overnight in BHI broth, centrifuged and concentrated 10-fold. Aliquots (0.1 ml) of each culture were plated onto pre-reduced BHI agar plates containing 0.32 μg/ml fidaxomicin or 0.5 μg/ml rifaximin, representing 4 x MICs. Similarly, WT and mutator cultures (>10 each) were concentrated and plated onto agar plates containing 2 or 4 x MIC (i.e. 0.5 or 1 μg/ml of metronidazole). After 48 hours of incubation, the mutation frequencies were calculated as the number of resistant colonies divided by total viable counts.

### Experimental evolution by serial passaging

From an overnight culture on agar, *C. difficile* colonies were resuspended into 1 ml of BHI broth to produce an optical density of OD_600_nm of ∼0.8-1.0. At each passage step, an aliquot of 0.01 ml (inocula of 10^5^-10^6^ CFU/mL) was spread onto BHI agars containing 0.25 to 64 × MIC of metronidazole. After 48 to 72 hours of incubation, colonies were isolated from the highest concentration permitting visible growth and were resuspended in fresh broth, before passaging the suspension onto higher drug concentrations. The remaining was stored in glycerol.

### Genome sequencing and analysis

Whole genome sequencing was done by paired end sequencing at SeqMatics LLC (California) and MRDNA (Texas). CLC Genomics Workbench version 12 (Qiagen) was used to *de novo* assemble WT ATCC 700057. The assembled genome of the WT was annotated using Rapid Annotation using Subsystem Technology (RAST) [49] and was used for mapping the mutator 700057Δ*mutSL* and metronidazole-resistant mutants (JWD-1, JWD-2, JWD-3 and JWD-4). Sequence variations were identified using “Quality-based variant detection” tool in CLC Genomics Workbench with default parameters (≥10-fold coverage of the reference position and sequence variation ≥35% of mapped reads). Mutations detected in the endpoint mutants, compared to the WT, were screened against the 700057Δ*mutSL* to remove shared variations, to focus on those that arose during the evolutions. For confirmation, the variations in ≥90% of mapped reads were further analyzed manually. The coverage for each genomic location was calculated in the CLC software and zero coverage regions were then identified using a customized perl script. Genetic changes were confirmed by Sanger sequencing.

### Growth kinetics of genetically manipulated strains

Two-fold serial dilutions of drugs was made in sterile pre-reduced BHI broth and 2% sterile xylose was used to induce antisense expression. Overnight grown cultures were diluted 1:100 and inocula (OD_600_ ∼0.3) was added to the pre-reduced 96 well plates and incubated in an anaerobic chamber for 24 hours. Plates were read for Absorbance_600_ using Synergy H1 Microplate Reader (BioTek).

### Quantification of intracellular iron by ICP-OES

Overnight grown cells were subcultured (1:100) in fresh BHI broth and harvested after 6 h for ICP-OES analysis to determine the cellular iron levels. Samples were washed twice, centrifuged at 5000 rpm and air dried. Samples were first digested in a microwave at 250°C at an initial pressure of 40 bar, at 250°C the pressure was about 80 bar. Precisely weighed samples were loaded into the digestion tube and 4 ml 16N HNO_3_ was added. After digestion, concentrated HNO_3_ was evaporated at 100°C, before addition of 5 ml 2% HNO_3_ and heating at 120°C to re-dissolve the samples. After weighing, iron content was measured with Agilent 725 ICP-OES. Calibration curves were made using standard solutions of iron at concentrations of 5 ppb-500 ppb.

### LC-MS/MS analysis of cellular lysates

Ten-fold concentrated, logarithmically growing cells (OD_600_ ∼0.3) were exposed to dimetridazole 1 mM and samples were harvested after 1 hour of incubation. Cellular lysates (10 μl) were mixed with methanol (90 μl), vortexed and centrifuged at 15,000 x g for 10 min. Supernatant (5 μl) was injected on to UHPLC-Q Exactive MS system for analysis. The concentration of dimetridazole was calculated based on the standard curve; further details are found in the supplementary.

### Analysis of DNA damage

Samples were harvested as above, after exposing to metronidazole 8 μg/ml and DNA damage was analyzed by alkaline agarose gel electrophoresis as described (48), except for the following changes. The lysis buffer contained lysozyme (0.5 mg/ml) and the agarose plugs were incubated at 37° C for 4 h. 0.5% v/v of hydrogen peroxide was a positive control to assess DNA fragmentation.

### Quantification of non-protein thiols

Samples harvested as above, were processed in 5% w/v trichloroacetic acid as previously described (38). Thiol content was then measured using the Thiol Fluorescent Detection Kit (Invitrogen), according to the manufacturer’s instructions.

### Transcriptional analysis

After harvesting the samples as above, bacterial RNAprotect™ reagent (QIAGEN) was then added to cultures and RNA isolated using Qiagen’s RNeasy mini kit. RNA was converted to cDNA by M-MLV reverse transcriptase (Quantabio) using qScript cDNA SuperMix. qScript One-Step SYBR Green qRT-PCR Kit, ROX (Quantabio) and gene-specific primers were used to amplify genes in Applied Biosystems ViiA7 real-time PCR machine. Transcript levels were calculated by the comparative Ct Method (ΔΔCT method) and data normalized to 16S rRNA.

### Quantification of lactate and pyruvate

Samples were harvested as above; lactate and pyruvate were detected using Pyruvate Colorimetric/Fluorometric Assay kit (Biovison) and Lactate-Glo™ Assay kit (Promega).

## ACKNOWLEDGEMENTS

This work was in part funded by grants R56AI126881 and R01AI139261 to J.G.H. from the National Institute of Allergy and Infectious Diseases at the National Institutes of Health. The funders had no role in study design, data collection and interpretation of the findings, or in the writing and submission of the manuscript.

ICP-OES analysis was conducted at the ICP lab, Department of Earth and Atmospheric Sciences, University of Houston, Texas. LC-MS/MS analysis was performed at NMR and Drug Metabolism core, Advanced Technology Core, Baylor College of Medicine, Houston, Texas.

## REFERENCES

1. Lessa FC, Mu Y, Bamberg WM, Beldavs ZG, Dumyati GK, Dunn JR, Farley MM, Holzbauer SM, Meek JI, Phipps EC, Wilson LE, Winston LG, Cohen JA, Limbago BM, Fridkin SK, Gerding DN, McDonald LC. 2015. Burden of *Clostridium difficile* infection in the United States. N Engl J Med 372:825–34.

2. Freeman J, Bauer MP, Baines SD, Corver J, Fawley WN, Goorhuis B, Kuijper EJ, Wilcox MH. 2010. The changing epidemiology of *Clostridium difficile* infections. Clin Microbiol Rev 23:529–49.

3. Pelaez T, Alcala L, Alonso R, Rodriguez-Creixems M, Garcia-Lechuz JM, Bouza E. 2002. Reassessment of *Clostridium difficile* susceptibility to metronidazole and vancomycin. Antimicrob Agents Chemother 46:1647–50.

4. Dingsdag SA, Hunter N. 2018. Metronidazole: an update on metabolism, structure-cytotoxicity and resistance mechanisms. J Antimicrob Chemother 73:265–279.

5. McDonald LC, Gerding DN, Johnson S, Bakken JS, Carroll KC, Coffin SE, Dubberke ER, Garey KW, Gould CV, Kelly C, Loo V, Shaklee Sammons J, Sandora TJ, Wilcox MH. 2018. Clinical Practice Guidelines for *Clostridium difficile* Infection in Adults and Children: 2017 Update by the Infectious Diseases Society of America (IDSA) and Society for Healthcare Epidemiology of America (SHEA). Clin Infect Dis 66:e1–e48.

6. Freeman J, Vernon J, Pilling S, Morris K, Nicholson S, Shearman S, Longshaw C, Wilcox MH, Pan-European Longitudinal Surveillance of Antibiotic Resistance among Prevalent Clostridium difficile Ribotypes Study G. 2018. The ClosER study: results from a three-year pan-European longitudinal surveillance of antibiotic resistance among prevalent *Clostridium difficile* ribotypes, 2011-2014. Clin Microbiol Infect 24:724–731.

7. Thorpe CM, McDermott LA, Tran MK, Chang J, Jenkins SG, Goldstein EJC, Patel R, Forbes BA, Johnson S, Gerding DN, Snydman DR. 2019. U.S.-Based National Surveillance for Fidaxomicin Susceptibility of *Clostridioides difficile*-Associated Diarrheal Isolates from 2013 to 2016. Antimicrob Agents Chemother 63.

8. Adler A, Miller-Roll T, Bradenstein R, Block C, Mendelson B, Parizade M, Paitan Y, Schwartz D, Peled N, Carmeli Y, Schwaber MJ. 2015. A national survey of the molecular epidemiology of *Clostridium difficile* in Israel: the dissemination of the ribotype 027 strain with reduced susceptibility to vancomycin and metronidazole. Diagn Microbiol Infect Dis 83:21–4.

9. Lynch T, Chong P, Zhang J, Hizon R, Du T, Graham MR, Beniac DR, Booth TF, Kibsey P, Miller M, Gravel D, Mulvey MR, Canadian Nosocomial Infection Surveillance P. 2013. Characterization of a stable, metronidazole-resistant *Clostridium difficile* clinical isolate. PLoS One 8:e53757.

10. Moura I, Monot M, Tani C, Spigaglia P, Barbanti F, Norais N, Dupuy B, Bouza E, Mastrantonio P. 2014. Multidisciplinary analysis of a nontoxigenic *Clostridium difficile* strain with stable resistance to metronidazole. Antimicrob Agents Chemother 58:4957–60.

11. Boekhoud IM, Hornung BVH, Sevilla E, Harmanus C, Bos-Sanders I, Terveer EM, Bolea R, Corver J, Kuijper EJ, Smits WK. 2020. Plasmid-mediated metronidazole resistance in *Clostridioides difficile*. Nat Commun 11:598.

12. Moura I, Spigaglia P, Barbanti F, Mastrantonio P. 2013. Analysis of metronidazole susceptibility in different *Clostridium difficile* PCR ribotypes. J Antimicrob Chemother 68:362–5.

13. Kumar M, Adhikari S, Hurdle JG. 2014. Action of nitroheterocyclic drugs against *Clostridium difficile*. Int J Antimicrob Agents 44:314–9.

14. Canfield GS, Schwingel JM, Foley MH, Vore KL, Boonanantanasarn K, Gill AL, Sutton MD, Gill SR. 2013. Evolution in fast forward: a potential role for mutators in accelerating *Staphylococcus aureus* pathoadaptation. J Bacteriol 195:615–28.

15. Long H, Miller SF, Strauss C, Zhao C, Cheng L, Ye Z, Griffin K, Te R, Lee H, Chen CC, Lynch M. 2016. Antibiotic treatment enhances the genome-wide mutation rate of target cells. Proc Natl Acad Sci U S A 113:E2498–505.

16. Lenhart JS, Pillon MC, Guarne A, Biteen JS, Simmons LA. 2015. Mismatch repair in Gram-positive bacteria. Res Microbiol doi:10.1016/j.resmic.2015.08.006.

17. Leeds JA, Sachdeva M, Mullin S, Barnes SW, Ruzin A. 2014. In vitro selection, via serial passage, of *Clostridium difficile* mutants with reduced susceptibility to fidaxomicin or vancomycin. J Antimicrob Chemother 69:41–4.

18. Zhou Y, Mao L, Yu J, Lin Q, Luo Y, Zhu X, Sun Z. 2019. Epidemiology of *Clostridium difficile* infection in hospitalized adults and the first isolation of *C. difficile* PCR ribotype 027 in central China. BMC Infect Dis 19:232.

19. Cartman ST, Kelly ML, Heeg D, Heap JT, Minton NP. 2012. Precise manipulation of the *Clostridium difficile* chromosome reveals a lack of association between the tcdC genotype and toxin production. Appl Environ Microbiol 78:4683–90.

20. Ho TD, Ellermeier CD. 2015. Ferric Uptake Regulator Fur Control of Putative Iron Acquisition Systems in *Clostridium difficile*. J Bacteriol 197:2930–40.

21. Hastie JL, Hanna PC, Carlson PE. 2018. Transcriptional response of *Clostridium difficile* to low iron conditions. Pathog Dis 76.

22. Yeom J, Imlay JA, Park W. 2010. Iron homeostasis affects antibiotic-mediated cell death in *Pseudomonas* species. J Biol Chem 285:22689–95.

23. Veeranagouda Y, Husain F, Boente R, Moore J, Smith CJ, Rocha ER, Patrick S, Wexler HM. 2014. Deficiency of the ferrous iron transporter FeoAB is linked with metronidazole resistance in *Bacteroides fragilis*. J Antimicrob Chemother 69:2634–43.

24. Pieulle L, Magro V, Hatchikian EC. 1997. Isolation and analysis of the gene encoding the pyruvate-ferredoxin oxidoreductase of *Desulfovibrio africanus*, production of the recombinant enzyme in *Escherichia coli*, and effect of carboxy-terminal deletions on its stability. J Bacteriol 179:5684–92.

25. Chen PY, Aman H, Can M, Ragsdale SW, Drennan CL. 2018. Binding site for coenzyme A revealed in the structure of pyruvate:ferredoxin oxidoreductase from *Moorella thermoacetica*. Proc Natl Acad Sci U S A 115:3846–3851.

26. Zuker M. 2003. Mfold web server for nucleic acid folding and hybridization prediction. Nucleic Acids Res 31:3406–15.

27. Digby WSaD. 1999. mRNAs have greater negative folding free energies than shuffled or codon choice randomized sequences. Nucl Acid Research 27:1578–1584.

28. Yeo WS, Lee JH, Lee KC, Roe JH. 2006. IscR acts as an activator in response to oxidative stress for the suf operon encoding Fe-S assembly proteins. Mol Microbiol 61:206–18.

29. Giel JL, Rodionov D, Liu M, Blattner FR, Kiley PJ. 2006. IscR-dependent gene expression links iron-sulphur cluster assembly to the control of O2-regulated genes in *Escherichia coli*. Mol Microbiol 60:1058–75.

30. Andre G, Haudecoeur E, Courtois E, Monot M, Dupuy B, Rodionov DA, Martin-Verstraete I. 2017. Cpe1786/IscR of *Clostridium perfringens* represses expression of genes involved in Fe-S cluster biogenesis. Res Microbiol 168:345–355.

31. Santos JA, Alonso-Garcia N, Macedo-Ribeiro S, Pereira PJ. 2014. The unique regulation of iron-sulfur cluster biogenesis in a Gram-positive bacterium. Proc Natl Acad Sci U S A 111:E2251–60.

32. Fleischhacker AS, Stubna A, Hsueh KL, Guo Y, Teter SJ, Rose JC, Brunold TC, Markley JL, Munck E, Kiley PJ. 2012. Characterization of the [2Fe-2S] cluster of *Escherichia coli* transcription factor IscR. Biochemistry 51:4453–62.

33. Pettersen EF, Goddard TD, Huang CC, Couch GS, Greenblatt DM, Meng EC, Ferrin TE. 2004. UCSF Chimera--a visualization system for exploratory research and analysis. J Comput Chem 25:1605–12.

34. Guerlesquin MBaFo. 1988. Structure, function and evolution of bacterial ferredoxins. FEMS Microbiology Reviews 54 155–176.

35. Carlier JP, Sellier N, Rager MN, Reysset G. 1997. Metabolism of a 5-nitroimidazole in susceptible and resistant isogenic strains of *Bacteroides fragilis*. Antimicrob Agents Chemother 41:1495–9.

36. Cudmore SL, Delgaty KL, Hayward-McClelland SF, Petrin DP, Garber GE. 2004. Treatment of infections caused by metronidazole-resistant *Trichomonas vaginalis*. Clin Microbiol Rev 17:783-93, table of contents.

37. Marreddy RKR, Wu X, Sapkota M, Prior AM, Jones JA, Sun D, Hevener KE, Hurdle JG. 2019. The Fatty Acid Synthesis Protein Enoyl-ACP Reductase II (FabK) is a Target for Narrow-Spectrum Antibacterials for *Clostridium difficile* Infection. ACS Infect Dis 5:208–217.

38. Leitsch D, Kolarich D, Wilson IB, Altmann F, Duchene M. 2007. Nitroimidazole action in *Entamoeba histolytica*: a central role for thioredoxin reductase. PLoS Biol 5:e211.

39. Leitsch D, Schlosser S, Burgess A, Duchene M. 2012. Nitroimidazole drugs vary in their mode of action in the human parasite *Giardia lamblia*. Int J Parasitol Drugs Drug Resist 2:166–70.

40. Huseby DL, Pietsch F, Brandis G, Garoff L, Tegehall A, Hughes D. 2017. Mutation Supply and Relative Fitness Shape the Genotypes of Ciprofloxacin-Resistant *Escherichia coli*. Mol Biol Evol 34:1029–1039.

41. de Freitas MC, Resende JA, Ferreira-Machado AB, Saji GD, de Vasconcelos AT, da Silva VL, Nicolas MF, Diniz CG. 2016. Exploratory Investigation of *Bacteroides fragilis* Transcriptional Response during *In vitro* Exposure to Subinhibitory Concentration of Metronidazole. Front Microbiol 7:1465.

42. Imlay J, Fridovich I. 1992. Exogenous quinones directly inhibit the respiratory NADH dehydrogenase in *Escherichia coli.* Arch Biochem Biophys 296:337–46.

43. Fuangthong M, Jittawuttipoka T, Wisitkamol R, Romsang A, Duang-nkern J, Vattanaviboon P, Mongkolsuk S. 2015. IscR plays a role in oxidative stress resistance and pathogenicity of a plant pathogen, *Xanthomonas campestris*. Microbiol Res 170:139–46.

44. Chong PM, Lynch T, McCorrister S, Kibsey P, Miller M, Gravel D, Westmacott GR, Mulvey MR, Canadian Nosocomial Infection Surveillance P. 2014. Proteomic analysis of a NAP1 *Clostridium difficile* clinical isolate resistant to metronidazole. PLoS One 9:e82622.

45. Choi SS, Chivers PT, Berg DE. 2011. Point mutations in *Helicobacter pylori*’s fur regulatory gene that alter resistance to metronidazole, a prodrug activated by chemical reduction. PLoS One 6:e18236.

46. Samarawickrema NA, Brown DM, Upcroft JA, Thammapalerd N, Upcroft P. 1997. Involvement of superoxide dismutase and pyruvate:ferredoxin oxidoreductase in mechanisms of metronidazole resistance in *Entamoeba histolytica*. J Antimicrob Chemother 40:833–40.

47. Tsugawa H, Suzuki H, Satoh K, Hirata K, Matsuzaki J, Saito Y, Suematsu M, Hibi T. 2011. Two amino acids mutation of ferric uptake regulator determines *Helicobacter pylori* resistance to metronidazole. Antioxid Redox Signal 14:15–23.

48. Zirkle RE, Krieg NR. 1996. Development of a method based on alkaline gel electrophoresis for estimation of oxidative damage to DNA in *Escherichia coli*. J Appl Bacteriol 81:133–8.

